# The presence of corticomuscular coherence during unipedal stance

**DOI:** 10.1101/2021.07.12.452017

**Authors:** Sylmina Dalily Alkaff, Junichi Ushiyama

## Abstract

Standing in unipedal stance requires higher effort to maintain posture balance within narrow base-of-support. Although changes in cortical activity are known to occur as standing task difficulty increased, it is unclear whether it indicates a change in cortical control of muscle activity. To elucidate this point, this study examined corticomuscular coherence (CMC) between the sensorimotor cortex and ankle joint muscles during three tasks such as bipedal stance, unipedal stance, and isometric contraction tasks. For twenty-one healthy participants, we measured the maximal peak of CMC (CMCmax) between electroencephalograms overlying the foot representation area and surface electromyograms from the tibialis anterior (TA), medial gastrocnemius (MG), lateral gastrocnemius (LG), and soleus (SOL), respectively, for each task. We also measured the center of pressure (COP) during both stance tasks. Although there was no significant CMC during bipedal stance in all participants, most participants (n = 14) showed significant CMC for all muscles during unipedal stance with larger COP fluctuation in most participants. Indeed, there was significant difference in CMCmax between unipedal and bipedal stance tasks. The greater CMC would indicate increased cortical control of muscle activity to compensate for postural instability during unipedal stance.

## 1. Introduction

Standing on one leg (unipedal stance) is performed in various daily activities, such as putting on pants, climbing stairs, and playing a range of sports. This stance challenges the ability to maintain postural balance with a high risk of falling due to narrow base-of-support (BOS), which is the enclosing contact area between feet and the supporting surface. Therefore, the unipedal stance is often implemented for fall risk assessment, balance training and rehabilitation programs ^1–4^.

Unipedal stance requires rapid sensorimotor integration to maintain the center of mass (COM) within the BOS. Thus, it is important to understand the neural mechanisms that control postural balance in this stance. Previous studies reported that electromyogram (EMG) activity, primarily in the ankle joints, was increased during unipedal stance with a larger center of pressure (COP) displacement in young adults ^5–7^. These findings suggest that increased muscle activity is required to compensate larger COP fluctuations caused by higher postural demand in unipedal stance. Additionally, increased theta-band power and decreased alpha-band power were observed in electroencephalogram (EEG) signals over the sensorimotor cortex as the standing task difficulty increased by changing the stance from bipedal to unipedal ^8^. Thus, it is reasonable to assume that the sensorimotor cortex is more involved in the modulation of ankle muscle activity during unipedal stance, to accommodate the increasing task difficulty. However, there has been no direct evidence supporting this assumption to date.

One approach for determining the involvement of the sensorimotor cortex in motor control is analyzing the functional connection between cortical and muscle activities. Coherence between EEG over the sensorimotor cortex and EMG from the contracting muscle, known as corticomuscular coherence (CMC), can provide such clarification. When performing isometric voluntary contraction, CMC is typically observed within the β-band (15–35 Hz) ^9–12^. However, only a few studies have investigated CMC modulation during standing. Masakado et al. ^13^ investigated CMC in several postural tasks and reported that significant β-band CMC was obtained only during foot stomping tasks. Jacobs et al. ^14^ reported changes in β-band CMC during bipedal stance depending on the stance width, surface, and vision. However, these studies measured EMGs from a limited number of ankle joint muscles. Because the functional roles played in postural control differ across synergistic and/or antagonistic muscles around the ankle joint, it is important to analyze these muscles simultaneously to obtain a more complete understanding of the mechanisms underlying postural control.

The aim of the present study was to compare the degree of cortical involvement in muscle activity to postural control between bipedal and unipedal stances by analyzing CMC. To this end, we recorded EMGs from four major muscles acting on the ankle joints of both legs (tibialis anterior [TA], medial gastrocnemius [MG], lateral gastrocnemius [LG], and soleus [SOL]), and measured their CMC with EEG over the sensorimotor cortex. We compared the CMC during both stance tasks with that during isometric contraction task.

## 2. Results

The results of the Waterloo Footedness Questionnaire-Revised (WFQ-R) questionnaire revealed that all participants were right leg dominant. Fourteen participants showed significant maximal peak value of CMC (CMCmax) in at least one muscle during the unipedal stance task.

### 2.1 Typical example of EEG, EMG, and COP raw data in standing tasks

To depict the difference between bipedal and unipedal stance tasks, we showed typical EEG and rectified EMG signals during each standing task from a participant (Figure 2). The MG and SOL muscles were active during both bipedal and unipedal stance tasks. The TA and LG muscles showed little activity during the bipedal stance task, while they showed increased activity during the unipedal stance task. Moreover, we observed clearer oscillations in all muscles during the unipedal stance task, while the MG and SOL showed tonic (rather than rhythmic) activations during the bipedal stance task.

**Figure 1.**
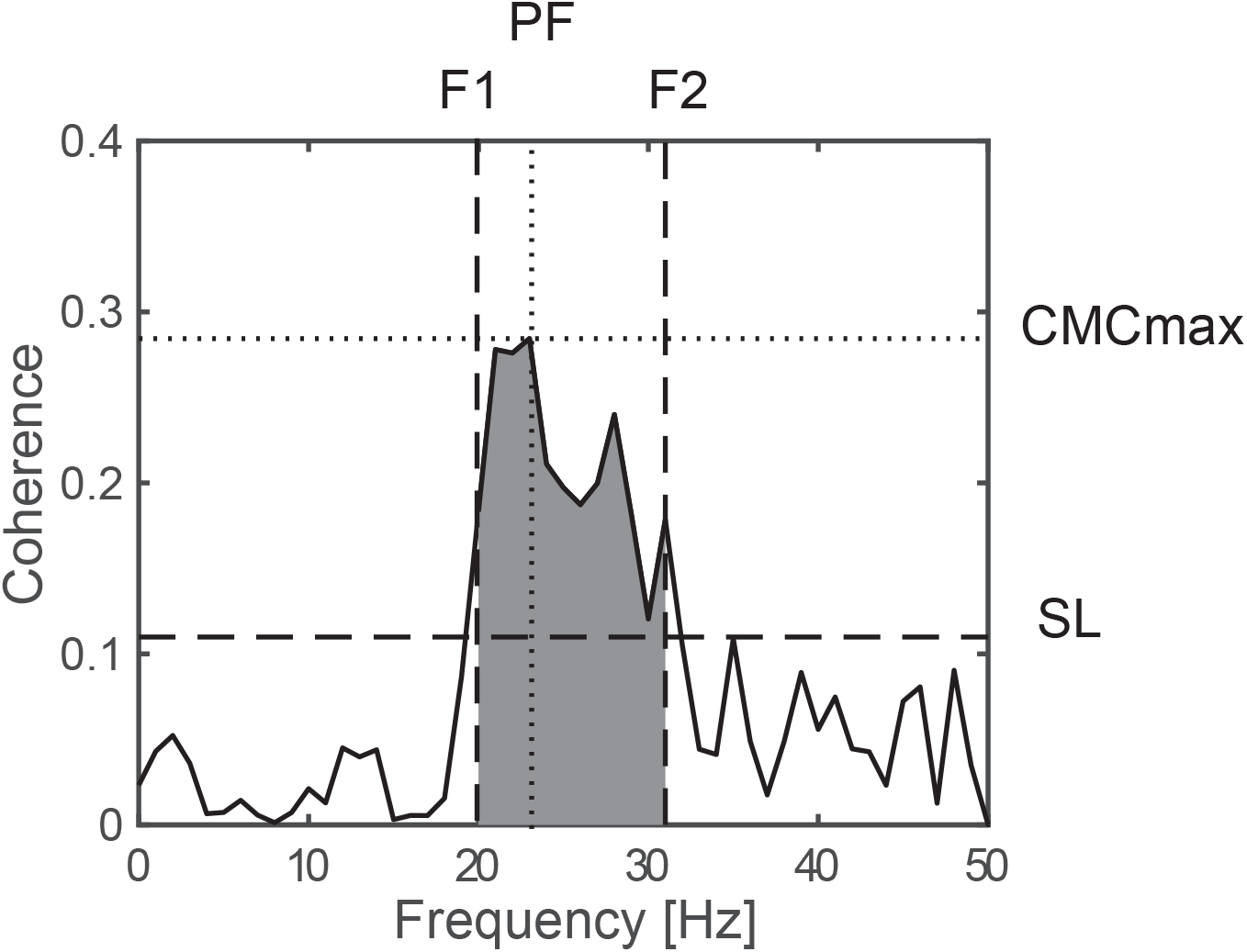
An example of a coherence spectrum between electroencephalogram (EEG) over the sensorimotor cortex and rectified electromyogram (EMG). The figure shows data from the tibialis anterior muscle (TA) during the isometric contraction task. We calculated the maximum value of CMC (CMCmax), the peak frequency at which CMCmax was observed (PF), the frequency at which coherence first met the significance level (SL) when traced backwards (F1) and forward (F2) from PF. CMCmax was considered statistically significant when it was greater than SL.

**Figure 2.**
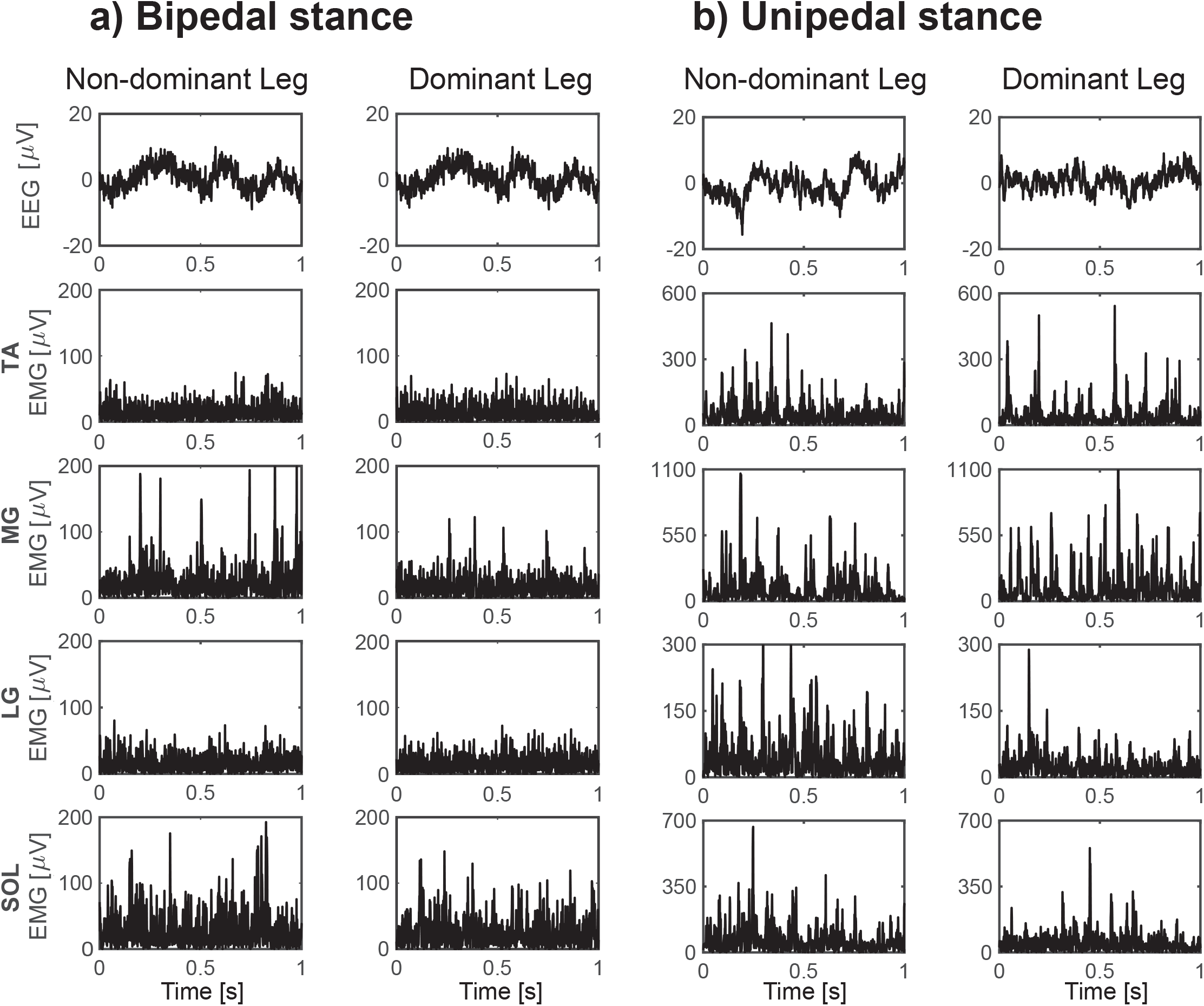
Examples of raw EEG over the sensorimotor cortex and rectified EMG from the four muscles (TA; Medial Gastrocnemius, MG; Lateral Gastrocnemius, LG; Soleus, Sol) on each leg during the bipedal stance (a) and unipedal stance (b) tasks. Data were taken from a participant who showed significant CMC only during the unipedal stance task. Note that the unipedal stance task data were collected separately from each leg, while the bipedal stance task data were collected simultaneously from both legs.

The results revealed an increase in COP fluctuation during both unipedal stance tasks compared with the bipedal stance task (Figure 3). This finding indicates that participants were least stable during both unipedal stance tasks than during the bipedal stance task.

**Figure 3.**
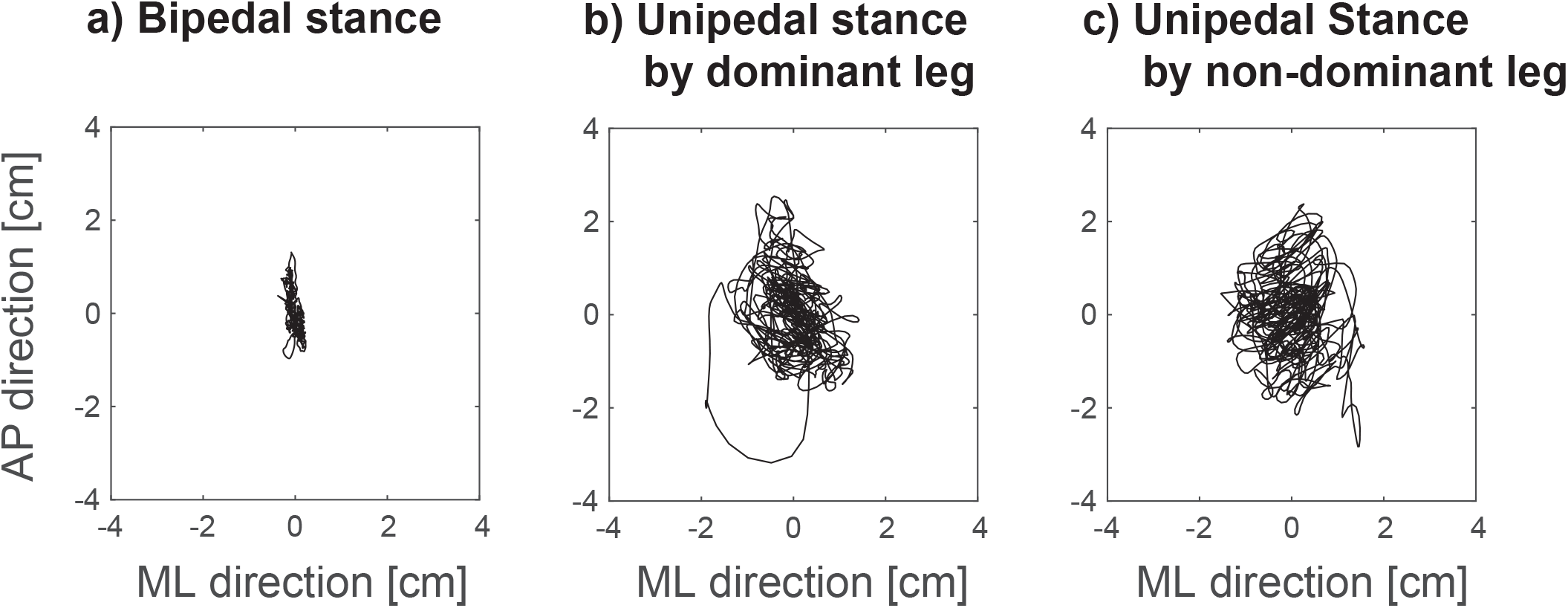
Typical examples of center of pressure (COP) displacement signals in anterior-posterior (AP) and medio-lateral (ML) directions from the same participant as in Figure 2. The figure shows data from the bipedal stance task (a), the unipedal stance task performed by the dominant leg (b), and the unipedal stance task performed by the non-dominant leg (c). During the unipedal stance task, COP fluctuation was larger compared with during the bipedal stance task.

### 2.2 Pooled CMC

To visualize the grand-average CMC spectra across participants, we showed pooled CMC spectra of each muscle in all tasks in Figure 4. The pooled CMC was not significant during bipedal stance task in all muscles. On the other hand, the pooled CMC was significantly observed in both unipedal stance and isometric contraction tasks in all muscles. Interestingly, although the significant pooled CMC was observed within the β-band (15-35 Hz) during isometric contraction task, it occurred mainly within the γ-band (36-50 Hz) during unipedal stance task. Additionally, significant CMC was also observed in lower frequency band (lower than the α-band) in some muscles during unipedal stance task. We suspect that this might be caused by EEG and EMG cable swaying due to increased body movement during unipedal stance.

**Figure 4.**
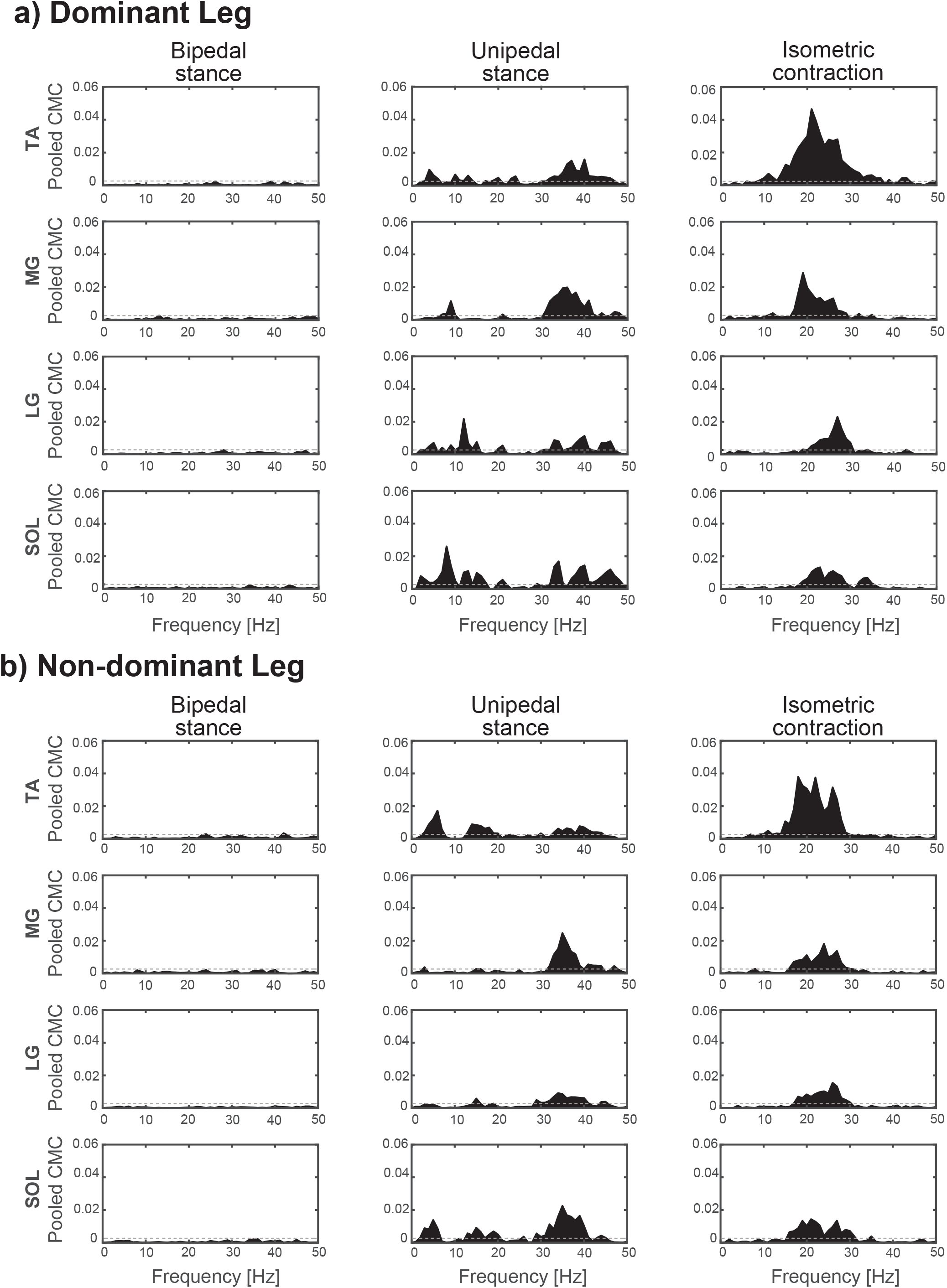
Effect of task on pooled CMC between EEG and EMG of four ankle muscles. Data from all participants were combined into one data to calculate pooled CMC of each muscle and task. Data are presented separately for each leg (**a**, dominant leg and **b**, non-dominant leg). In the coherence spectra, black lines indicate the calculated pooled CMC. SL is represented in horizontal dashed line.

### 2.3 Differences in COP parameters between bipedal and unipedal standing tasks

Figure 5 represents the group data from all COP parameters on each standing task, which are mean distance (MDIST), standard deviation (SD), mean velocity (MVELO), and mean frequency (MFREQ). One-way repeated measures ANOVA test revealed significant differences between tasks on all COP parameters (MDIST, F_2,26_ = 63.159, p < 0.001; SD, F_2,26_ = 63.962, p < 0.001; MVELO, F_2,26_ = 119.415, p < 0.001; MFREQ, F_2,26_ = 26.667; p < 0.001). Post-hoc analysis confirmed that all parameters during both unipedal stance tasks with the dominant leg (MDIST, 0.8425 ± 0.132 cm; SD, 0.966 ± 0.148 cm; MVELO, 1.594 ± 0.357 cm/s; MFREQ, 0.306 ± 0.065 Hz) and with the non-dominant leg (MDIST, 0.774 ± 0.133 cm; SD, 0.882 ± 0.151 cm; MVELO, 1.642 ± 0.390 cm/s; MFREQ, 0.344 ± 0.072 Hz) were significantly higher than those during the bipedal stance task (MDIST, 0.374 ± 0.104 cm; SD, 0.442 ± 0.118 cm; MVELO, 0.449 ± 0.071 cm/s; MFREQ, 0.203 ± 0.039 Hz) (all, p < 0.001). This finding indicates that humans are inherently unstable during unipedal stance. In addition, post-hoc analysis revealed that there was no significant difference in COP parameters between unipedal stance using the dominant and non-dominant legs, indicating that leg dominance did not affect instability of participants (MDIST, p = 0.520; SD, p = 0.418; MVELO, p = 0.899; MFREQ, p = 0.302).

**Figure 5.**
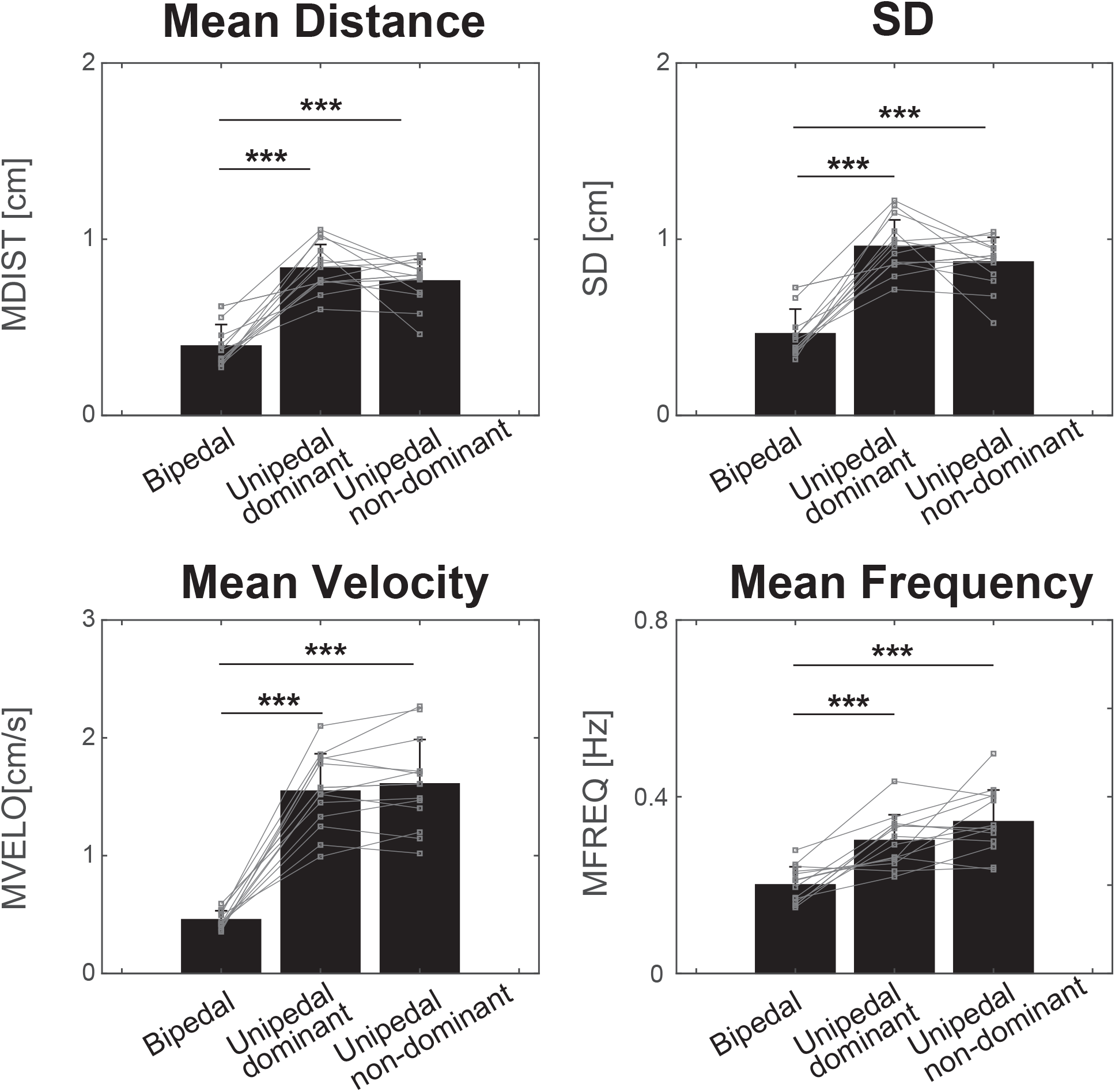
Group data (mean ± SD) for COP parameters on three tasks (bipedal stance, unipedal stance performed by the dominant leg, and unipedal stance performed by the non-dominant leg). Black bars indicate the average COP parameter value while the black line on the bar indicates standard deviation (SD) across participants. Individual data of each COP parameters are represented in grey lines. Significant differences between tasks are denoted as follows: *p < 0.05, **p < 0.01, ***p < 0.001.

### 2.4 Differences in CMC and EMG activity between standing and isometric contraction tasks

Figure 6 shows the group data for CMCmax from all muscles on each task. Two-way repeated measures ANOVA revealed that the main effect of task on CMCmax was significant for all muscles (TA, F_1.17,15.17_ = 14.419, p = 0.001; MG, F_1.24,16.1_ = 7.733, p = 0.01; LG, F_2,26_ = 5.876, p = 0.008; SOL, F_2,26_ = 4.528, p = 0.021). However, there was no significant effect of leg dominance (TA, F_1,13_ = 0.135, p = 0.719; MG, F_1,13_ = 3.741, p = 0.075; LG, F_1,13_ = 0.100, p = 0.757; SOL, F_1,13_ = 0.000, p = 0.986) and no significant interaction between leg dominance and task for all muscles (TA, F_1.35,17.53_ = 0.161, p = 0.767; MG, F_1.24,16.07_ = 0.875, p = 0.386; LG, F_2,26_ = 1.657, p = 0.210; SOL, F_1.40,18.27_ = 0.048, p = 0.902). For further statistical analysis, CMCmax data from both legs were combined and post-hoc tests were performed for comparing differences in CMCmax among tasks for each muscle. The results revealed that all muscles showed significantly higher CMCmax during the unipedal stance task (TA, 0.069 ± 0.033; MG, 0.060 ± 0.034; LG, 0.067 ± 0.030; SOL, 0.074 ± 0.038) compared with the bipedal stance task (TA, 0.037 ± 0.011; MG, 0.033 ± 0.011; LG, 0.037 ± 0.011; SOL, 0.041 ± 0.015) (TA, p = 0.002; MG, p = 0.016; LG, p = 0.003; SOL, p = 0.009).

**Figure 6.**
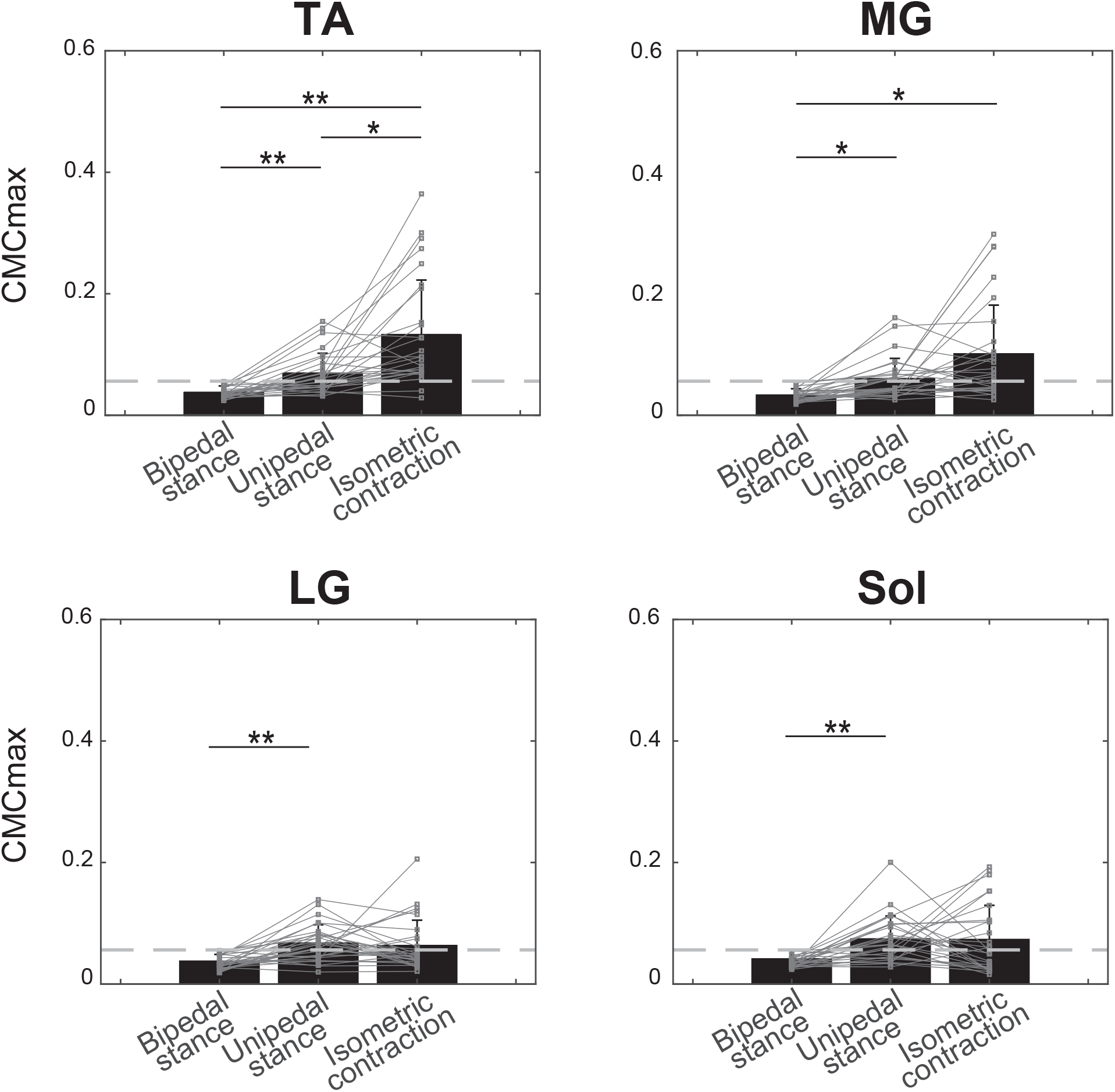
Group data (mean ± SD) for CMCmax from four muscles on three tasks (bipedal stance, unipedal stance, and isometric contraction). Note that the data from the dominant and non-dominant legs were combined into one figure for each muscle as the statistical analysis showed that there was no significant effect of leg dominance on CMCmax. Black bars indicate average CMCmax, while black lines on the bar indicate standard deviation (SD) across participants. Grey lines indicated individual CMCmax data. The SL is represented as horizontal dashed lines. CMCmax was higher during the unipedal and isometric contraction tasks compared with the bipedal stance task. Significant differences between tasks are denoted as follows: *p < 0.05, **p < 0.01, ***p < 0.001.

In addition, both the TA and MG muscles showed significantly higher CMCmax during the isometric contraction task (TA, 0.132 ± 0.090; MG, 0.101 ± 0.080) compared with the bipedal stance task (TA, p = 0.003; MG, p = 0.015). Only the TA muscle showed significantly higher CMC during the isometric contraction task compared with the unipedal stance task (p = 0.028). Although we could not perform statistical comparison of CMCmax frequency distribution between the unipedal stance and isometric contraction tasks because of inconsistency of numbers of data-points with significant CMCmax between tasks, we found a tendency for the peak frequency (PF) to shift to high β- or γ-band during the unipedal stance task in all muscles (Figure 7), similar to the pooled CMC result.

**Figure 7.**
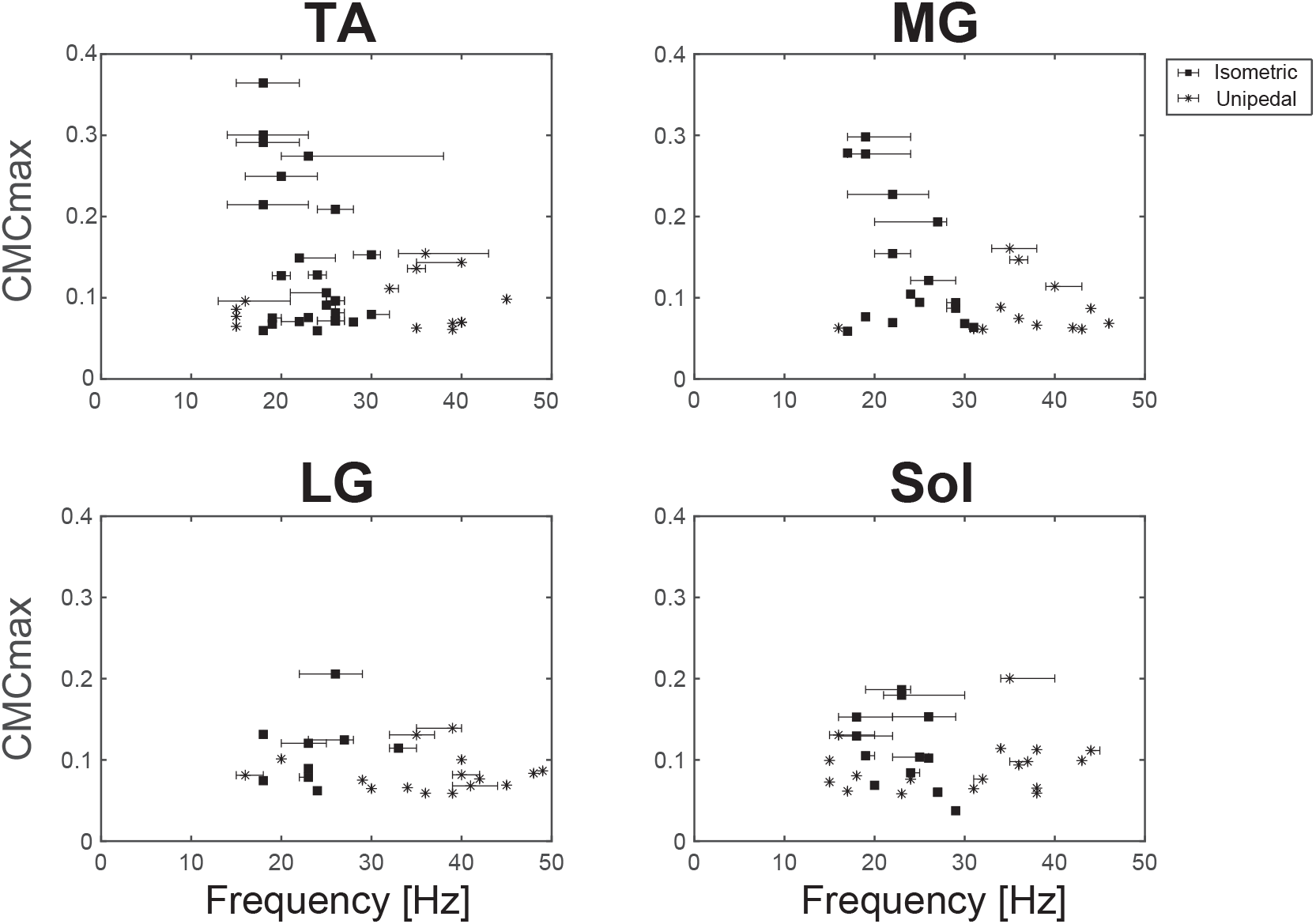
Distribution relationship between frequency width of significant CMC and CMCmax during isometric contraction and unipedal stance tasks. Data from dominant and non-dominant legs were merged into one figure for each muscle. Black squares indicate the PF value during the isometric contraction task, while asterisks indicate the PF value during the unipedal stance task. The error bar indicates the width of F1–F2.

Figure 8 shows the group data for mean EMG activity level from all muscles between standing tasks. Mean EMG activity level during the standing task was normalized to mean EMG activity level during maximum voluntary contraction (EMGmax). Two-way repeated measures ANOVA revealed that the main effect of task on the mean EMG activity level in all muscles was significant (TA, F_1,13_ = 37.014, p < 0.001; MG, F_1,13_ = 35.525, p < 0.001; LG, F_1,13_ = 16.850, p = 0.001; SOL, F_1,13_ = 18.939, p < 0.001). However, there was no significant effect of leg dominance (TA, F_1,13_ = 2.046, p = 0.176; MG, F_1,13_ = 0.544, p = 0.474; LG, F_1,13_ = 4.614, p = 0.051, SOL, F_1,13_ = 3.455, p = 0.086) and no significant interaction between leg dominance and task for all muscles (TA, F_1,13_ = 0.148, p = 0.707; MG, F_1,13_ = 0.282, p = 0.683; LG, F_1,13_ = 0.993, p = 0.337; SOL, F_1,13_ = 0.159, p = 0.697). Similar to CMCmax, mean EMG activity level data from both legs were combined and post-hoc tests were performed to compare the difference among tasks for each muscle. The results revealed that all muscles exhibited significantly higher mean EMG activity level in the unipedal stance task (TA, 11.031 ± 5.547 %EMGmax; MG, 29.746 ± 15.668 %EMGmax; LG, 18.503 ± 11.293 %EMGmax; SOL, 35.618 ± 16.774 %EMGmax) compared with the bipedal stance task (TA, 5.166 ± 3.945 %EMGmax; MG, 7.821 ± 5.231 %EMGmax; LG, 11.264 ± 6.802 %EMGmax; SOL, 22.986 ± 13.570 %EMGmax) (TA, p < 0.001; MG, p < 0.001; LG, p = 0.001; SOL, p < 0.001).

**Figure 8.**
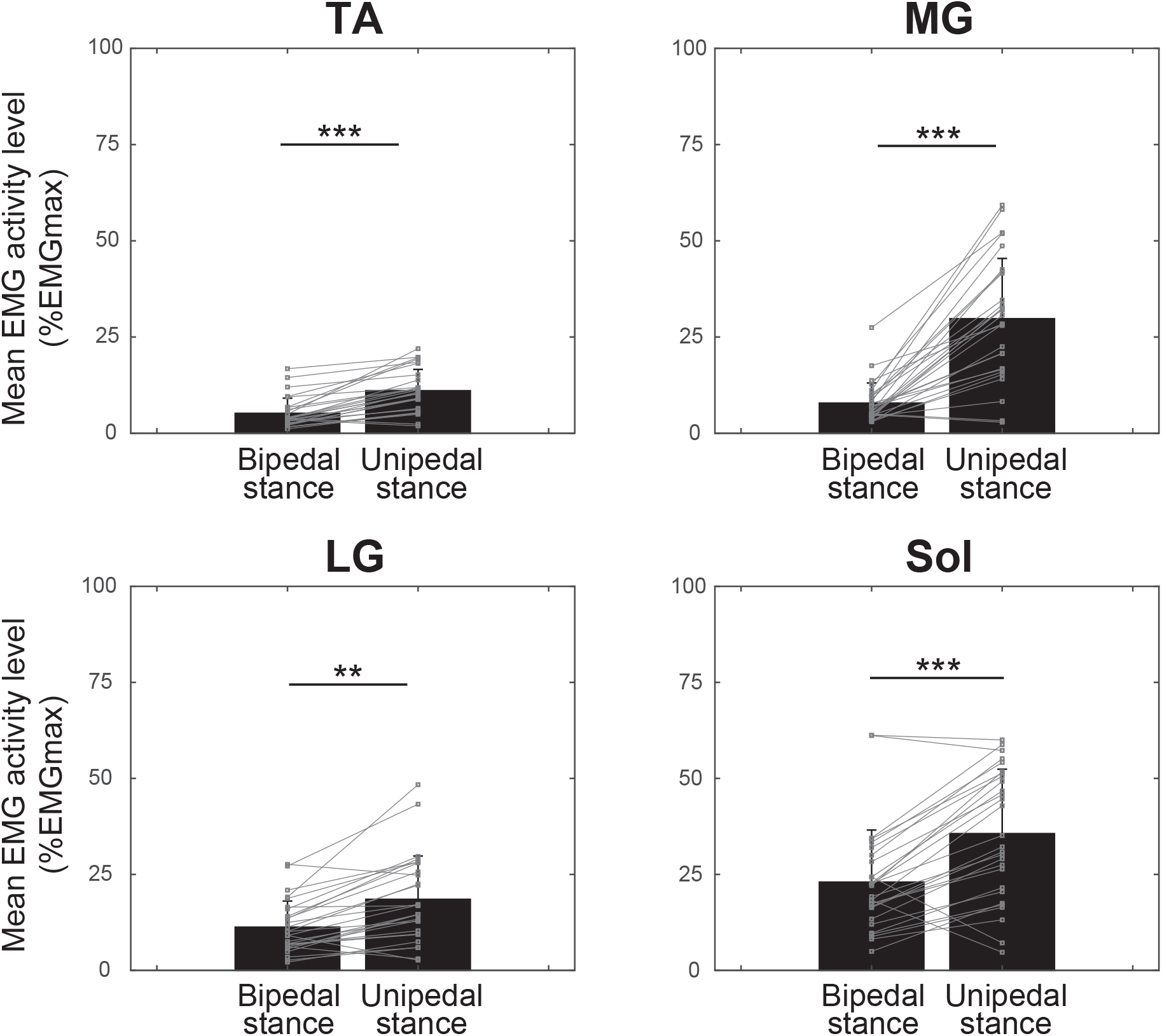
Group data (mean ± SD) for mean EMG activity level from four muscles on each task. Data from both dominant and non-dominant legs were merged into one figure for each muscle as the statistical analysis showed that there was no significant effect of leg dominance on mean EMG activity level. Black bars indicate the average value of mean EMG activity level while the black line on the bar indicates SD across participants. Individual EMG activity level data are represented in grey lines. Significant differences between tasks are denoted as follows: *p < 0.05, **p < 0.01, ***p < 0.001.

## 3. Discussion

The present study compared the degree of cortical involvement in postural control between bipedal and unipedal stance tasks, by measuring CMC between EEG over the sensorimotor cortex and EMG from ankle extensor and flexor muscles. In most participants, we found significant CMC for all muscles during the unipedal stance task with larger COP fluctuation, whereas no significant CMC was observed during the bipedal stance task.

### 3.1 Stronger CMC during unipedal stance indicated increased direct cortical involvement in postural control

The current findings are in accordance with previous reports ^13^, revealing no significant CMC from any muscles during bipedal stance. Because postural balance is regarded to be mainly regulated by the cerebellum ^30^, basal ganglia, and brainstem ^31^, less or no involvement of the sensorimotor cortex in postural balance control during bipedal stance would be expected to result in a lack of significant CMC in the bipedal stance task. However, in the present study, significant CMC was observed during the unipedal stance in most participants. Indeed, CMCmax was significantly stronger in the unipedal stance task compared with the bipedal stance task for all muscles among participants with significant CMC. Therefore, the degree of cortical involvement in the control of muscle activity around the ankle joint would increase during unipedal stance.

Biomechanically, the human body during standing can be represented as an inverted pendulum rotating around the ankle joint with COM located in front of the joint ^32^. During bipedal stance, postural sway in response to counteracting gravitational force occurred mainly in the AP direction and was found to be coherent with ankle extensor muscle activity ^33,34^. However, as COP fluctuation becomes larger and faster because of narrow BOS, postural control strategies would be expected to be modified during unipedal stance.

As a biarticular muscle crossing the ankle and knee joints, motor unit recruitment in the MG is known to be triggered by forward sway occurring during bipedal stance, resulting in a phasic muscular activity pattern ^35^. Consequently, MG muscle activity was reported to be coherent with body angle during bipedal stance ^36^. This suggests that MG is responsible for regulating body fluctuation in postural control. Regarding unipedal stance, greater effort would be expected to be required for MG to compensate larger body fluctuation occurring because of narrow BOS, as reflected by the increased muscle activity level. Increased CMC during the unipedal stance suggests that MG requires increased cortical involvement to regulate body fluctuation.

As a monoarticular muscle, SOL muscle activity was found to be linearly correlated with ankle angle and ankle torque during bipedal stance ^37,38^. This suggests that SOL is responsible for maintaining ankle joint tone during quiet standing. In the case of unipedal stance, narrow BOS caused destabilization of postural balance, requiring the SOL muscle to increase ankle joint tone. Increased CMC during unipedal stance suggests that the SOL muscle requires increased cortical involvement to increase ankle joint tone.

Research on the functional role of the LG muscle in postural control has been less extensive compared with the MG and SOL muscles ^33,39,40^. This is because postural sway during the bipedal stance occurs mainly in the AP direction, resulting in less contribution from the LG muscle to postural control. However, during unipedal stance, the BOS area was significantly decreased in the frontal plane, resulting in imbalance in ML postural sway. Significant CMC observed during unipedal stance indicated that increased LG muscle activity level required increased cortical involvement to maintain postural balance in the ML direction.

As an antagonist muscle from the ankle extensor muscles, TA muscle activation is not required during the bipedal stance because the focus of postural control during the bipedal stance is to counteract forward sway caused by gravitational pull. However, the contribution of TA muscle activity to postural control is known to be increased when greater backward sway occurs, either because of narrow BOS ^41,42^ or backward external perturbation ^43^. In the current study, CMC obtained from the TA muscle was increased during unipedal stance, where BOS is relatively narrow. This indicates that increased TA muscle activation requires increased cortical involvement to provide forward sway to prevent backward falling.

As such, each ankle joint muscle would be expected to play different functional roles for postural control depending on the requirement from the postural response. During bipedal stance, the postural response was sufficient to maintain postural balance within BOS with less to no contribution from the sensorimotor cortex to postural control. However, the decreased BOS area when standing in the unipedal stance induces an imbalance in postural sway in both the AP and ML directions, as reflected by larger and faster COP fluctuation. To compensate for the greater postural demand during the unipedal stance, each muscle must increase the effort to maintain postural balance by increasing the activity level, requiring additional involvement from the sensorimotor cortex to provide more voluntary postural adjustment.

### 3.2 CMC peak frequency tends to shift towards higher β-band or γ-band during unipedal stance

Pooled CMC results across participants revealed a tendency for significant CMC during unipedal stance to be observed within higher frequency bands for all muscles. Previous studies also reported a frequency shift toward the γ-band during a dynamic contraction task ^44,45^ or during stronger intensity of an isometric contraction task ^46–48^.

However, it should be noted that a frequency shift toward high β-band or γ-band during unipedal stance occurred in the present study despite the low to moderate muscle activity level (less than 60% EMGmax). Masakado et al. ^13^ also reported a similar frequency shift toward higher frequency bands during unipedal stance in one participant with significant CMC, despite all of the other participants showing no significant CMC. A possible explanation for this tendency is that larger and faster COP fluctuation during unipedal stance demands faster postural correction, thus requiring the sensorimotor system to integrate the relevant sensory information with motor commands at a higher rate. By increasing oscillation frequency (i.e., γ-band), more and faster information could be exchanged within the corticospinal loop. Therefore, frequency shift tendency in CMC may have occurred to facilitate the demand of rapid sensorimotor integration because of the difficulty in maintaining postural balance caused by narrow BOS in the unipedal stance task.

### 3.3 Technical limitations of the study

Several potential limitations of the current study should be considered. First, the presence of an anatomical buffer, such as skull thickness, cerebrospinal fluid, and location and orientation of the relevant cortical neuron, may have influenced the signals measured by scalp EEG ^49–51^. Similar anatomical issues, such as orientation, size, and depth of muscle fibers, can also affect surface EMG recordings ^52,53^. However, while these limitations might affect the individual amplitude of EEG and EMG signals, CMC is a measure of the constancy of the amplitude ratio and phase difference between two signals ^26^. Therefore, we believe that these technical limitations were not likely to have had a substantial influence on the present results.

Second, the present study focused only on muscles around the ankle joint. However, it is possible that other proximal muscles (i.e., muscles acting on the hip and trunk joint) could have contributed to postural control, particularly during unipedal stance. Older adults, for example, were found to exhibit increased hip angle movement accompanied by greater hip muscle EMG activity compared with younger adults during unipedal stance ^6,7^. If a similar study was expanded to include older adults, it would be necessary to record EMG signals from proximal muscles and evaluate CMC. However, younger adults are known to mainly adopt ankle strategies for postural control with little contribution from the hip joint during both bipedal and unipedal stance ^5–7^. Therefore, although the contribution of hip movements to postural control cannot be ignored, we believe that the present approach, which focused on CMC in ankle muscles, was effective for elucidating the neural mechanisms involved in postural control in young adults.

Finally, it is known that individual differences exist within CMC ^11,12^, suggesting that some individuals or populations may not exhibit significant CMC in any task. Although we rejected participants who did not show significant CMC for all tasks in the current study, this does not imply that they do not utilize neural interactions between the sensorimotor cortex and contracting muscle during the task. Further studies using other measurements would be needed if participants who did not show significant CMC were included.

### 3.4 Conclusion

The degree of cortical involvement in postural control during bipedal and unipedal stance was investigated by measuring CMC in muscles around the ankle joint. We found significant CMC during unipedal stance for all muscles, while no significant CMC was observed in the bipedal stance task. These findings suggest that more voluntary control of muscle activity is required to compensate for postural instability during unipedal stance.

## 4. Methods

### 4.1 Ethical Approval

The experiment protocols and procedures were conducted following the Declaration of Helsinki and were approved by the ethical committee of the Faculty of Policy Management, Faculty of Environment and Information Studies, and Graduate School of Media and Governance, Keio University (receipt number 167). All participants received a detailed explanation of the experiment and provided informed consent prior to data collection.

### 4.2 Participants

Twenty-one healthy young adults (thirteen male and eight female, aged 18–25 years) participated in this study. All participants reported no history of neuromuscular and/or musculoskeletal disorders. The dominant foot of each participant was determined using three sets of questions taken from WFQ-R questionnaire ^15^.

### 4.3 Experimental Protocol

Participants performed an isometric contraction task and two standing tasks (i.e., bipedal and unipedal stance tasks). In the bipedal stance task, participants were instructed to stand with their feet separated by 10 cm and to keep their arms at their sides. Foot positions were marked with white tape to ensure that the same position was maintained throughout measurements. In the unipedal stance task, participants were instructed to stand on one leg and keep their arms at their sides, in the same manner as in the bipedal stance task. The unipedal stance task was conducted with each leg. Both standing tasks required participants to maintain an upright position and focus their eyes on a fixed target on a screen positioned approximately 2 m in front of them. When participants were unable to keep their posture straight, the task was stopped and repeated after a brief rest of approximately 5 minutes. Two trials were conducted for each task, with a duration of approximately 70 s for each trial. The order of the tasks was randomized for each participant.

The isometric contraction task was performed approximately 5 minutes after the standing tasks. Participants sat on a chair with each foot fitted to an ankle dynamometer by attached straps. Participants’ knee joints were fully extended, while the ankle joints were set in a neutral position. Before the isometric contraction task, participants performed maximal voluntary contraction (MVC) of ankle dorsiflexion and plantarflexion with each leg for 3 s while remaining as stable as possible. We determined the MVC value as the peak torque from this measurement. After a sufficient rest period of approximately 60 s, participants performed isometric ankle dorsiflexion and plantarflexion on 10% MVC for each leg. Each contraction was performed for approximately 70 s, and repeated twice. The intensity level for this task was determined from the average EMG activity level from all muscles while performing a bipedal stance task in preliminary experiments. Visual feedback was provided on a display placed approximately 2 m in front of the participants, consisting of target torque level (10% MVC) represented by a blue line and an exerted torque level represented by a red line. Participants were instructed to keep the red line as closely as possible to the blue line.

### 4.4 Data Collection

We recorded scalp EEG signals over the foot representation area of the sensorimotor cortex. Five passive Ag/AgCl electrodes with a diameter of 18 mm were placed at Cz and its surrounding positions (FCz, CPz, C1, and C2) defined by the international 10-20 system on an EEG cap (g.GAMMAcap 1027; Guger Technologies, Graz, Austria). Reference and ground electrodes were placed on the right and left earlobes, respectively. Surface EMG signals were recorded from TA, MG, LG, and SOL muscles from both legs. Two passive bipolar Ag/AgCl disposable electrodes with a diameter of 10 mm were placed over each muscle belly with an inter-electrode distance of 20 mm. EMG ground electrodes were placed on the kneecap on the right leg of the participant. Participants’ skin was gently abrased and cleaned with alcohol before sensor placement. The torque signal measured during isometric contraction task was recorded using an ankle dynamometer (TCF100N; Takei Scientific Instruments Co., Ltd., Niigata, Japan).

Both EEG and EMG signals were amplified and band-passed filtered with a cut-off frequency of 0.5–100 Hz for EEG and 2–1000 Hz for EMG, using a 16-channel analog biosignal amplifier (g.BSamp 0201a; Guger Technologies, Graz, Austria). The torque signal was low-pass filtered using a second-order Butterworth filter at a cut-off frequency of 50 Hz and amplified using an amplifier (Kyowa Electronics Instruments Co., Ltd., Tokyo, Japan). All signals were then converted to digital signals at a sample rate of 1000 Hz by an analog-to-digital converter with 16-bit resolution (NI USB-6259 BNC, National Instruments, Austin, TX, USA) controlled by data-logger software using MATLAB (The Mathworks, Natick, MA, USA).

The COP signals were recorded using a Wii Balance Board (RVL-021; Nintendo, Kyoto, Japan). Previous studies reported that Wii Balance Board provided a reliable, cost-effective alternative to an industrial force plate ^16–19^. Signals were collected via Bluetooth at a sample rate of 30 Hz by another data-logger software program using MATLAB.

### 4.5 Data Analyses

The EEG signal from Cz was first Laplacian-filtered by subtracting the average potential from surrounding electrodes, then filtered again using a second-order Butterworth low-pass filter at 50 Hz ^20^. The EMG signal was full-wave rectified because grouped discharge patterns could be detected, and was thus suitable for CMC analysis ^21–23^.

The last 60 s period of each trial was extracted from EEG and EMG signals and combined into one 120 s data-set for further analysis. Prior to coherence analysis, both EEG and rectified EMG signals from each muscle were segmented into 1 s data windows, resulting in 120 data epochs without overlap. Each data epoch was then Hanning-windowed to reduce spectral leakage ^9,24,25^ before we calculated the coherence between EEG and EMG of each signal. Coherence was calculated using the following formula ^26^:

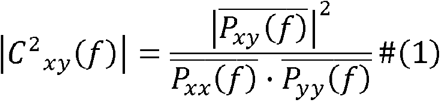

where 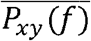 is the averaged cross-spectral density between EEG and rectified EMG signals, and 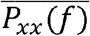 and 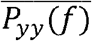 are the averaged power spectral densities of EEG and rectified EMG signals, respectively, at any given frequency *f*.

The calculated coherence value is between 0 and 1, where 1 indicates that two signals are perfectly correlated. A 95% confidence limit was determined as the SL of the coherence and the frequency range of interest was set between 3 and 50 Hz. In accordance with our previous studies (e.g., Ushiyama et al., 2017, 2011), we applied Bonferroni correction to the equation determining a 95% confidence limit to eliminate potential statistical errors caused by multiple comparisons, resulting in the following equation:

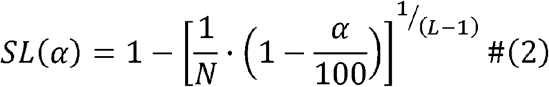

where *L* is the number of data segments, *N* is the number of frequency bins, and α is the 95% confidence limit. As *L* and *N* were set as 120 and 48, respectively, the SL in this study was determined to be 0.0565. The calculated coherence was considered to be significant when the value exceeded the SL.

To visualize the grand-average CMC spectra across all participants for different tasks, we also calculated pooled CMC of each muscle during each task. We first combined data from all participants and performed coherence calculation using the same method as the individual coherence.

In this study, we calculated CMCmax and PF within the β-band or γ-band. If the observed CMCmax value was above SL, we also calculated the frequencies where the coherence curve first met the SL when traced backward (F1) and forward (F2) from the PF ^27^, as shown in Figure 1. Among all participants who showed significant CMCmax, the PF was observed within either band depending on the task. We examined the mean EMG activity level from each muscle during all tasks. We first calculated the average value of rectified EMG within a 1 s window with maximal averaged torque output during MVC trials (EMGmax). We then calculated the average value of rectified EMG data for 120 s during each task and normalized it by the above-mentioned EMGmax.

COP data in the anterior-posterior (AP) and medio-lateral (ML) direction signals were low-pass filtered at 15 Hz with a fourth-order Butterworth filter. The last 60 s period of each trial was then extracted for further analysis. Based on previous studies, we subtracted the mean value from COP signals to remove the offset ^28,29^. We also calculated the resultant distance (RD) data series as the vector distance from mean COP to each data pair point in AP and ML data series, using the following equation:

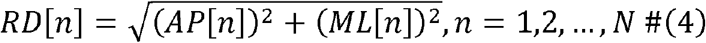

where *N* is the length of COP data included for the analysis. Various COP parameters can be used as measures of postural steadiness. We decided to calculate several parameters that have been most often used in posture and balance studies, such as SD, MVELO, MDIST, and for the RD direction. We chose to only include COP parameters in the RD direction as the result of COP fluctuation because the obtained results were similar to those in both the AP and ML directions. COP parameter results from each trial were then averaged for statistical analysis.

### 4.6 Statistics

Only participants showing significant CMC in at least one muscle during standing tasks were included for further analysis. To analyze the effects of standing tasks on each COP parameter in RD direction, we performed one-way repeated measures analysis of variance (ANOVA) to compare the average values of each COP parameter among stance tasks (i.e., bipedal stance, unipedal stance with dominant leg, and unipedal stance with the non-dominant leg). Two-way repeated measures ANOVA was performed to analyze the effects of leg dominance (dominant and non-dominant leg) and task (bipedal stance, unipedal stance, and isometric contraction tasks) on CMCmax obtained from each muscle. In addition, two-way repeated measures ANOVA was performed to determine whether leg dominance and task (bipedal and unipedal stance) also had an effect on mean EMG activity level from each muscle. Greenhouse-Geisser correction was applied for sphericity. If a significant interaction was obtained, one-way repeated measures ANOVA was performed to determine whether the effect of tasks was significant on each muscle from each leg. If the result was significant, post-hoc pairwise comparisons with Bonferroni corrections were performed to determine whether there was a significant difference between tasks. However, if the interaction was not significant and at least one of the factors was significant, post-hoc pairwise comparisons with Bonferroni correction were executed to determine whether there was a significant difference within the factor for each muscle. The significance level was set at α = 0.05 in all statistical tests. All statistical analyses were performed using SPSS software (IBM SPSS Statistics 26, IBM Developer Works, Tokyo, Japan).

## Data Availability

Data analyzed during this study are included in the Supplementary Information files of this article. Further data used to support the findings of this study are available from the corresponding author upon reasonable request.

## Acknowledgments

This work was supported by grants from the Grant-in-Aid for Scientific Research (B) (Japan Society for the Promotion of Science, JSPS) (grant number 20H04091) to JU, and a designated donation from Living Platform, Ltd, Japan to JU. We thank Ms. Tomomi Hamaoka, Ms. Kana Iijima, and Ms. Chieko Matsuda for their secretarial assistance. We thank Benjamin Knight, MSc., from Edanz (https://jp.edanz.com/ac) for editing a draft of this manuscript.

## Author Contributions

S.D.A. and J.U. conceptualized and designed the study, interpreted data, wrote the manuscripts. S.D.A. acquired and analyzed data. J.U. acquired funding and supervised the study.

## References

1. Hurvitz, E. A., Richardson, J. K., Werner, R. A., Ruhl, A. M. & Dixon, M. R. Unipedal stance testing as an indicator of fall risk among older outpatients. Arch. Phys. Med. Rehabil. 81, 587–591 (2000).

2. Trojian, T. H. & McKeag, D. B. Single leg balance test to identify risk of ankle sprains. Br. J. Sports Med. 40, 610–613 (2006).

3. Zech, A. et al. Neuromuscular training for rehabilitation of sports injuries: A systematic review. Med. Sci. Sports Exerc. 41, 1831–1841 (2009).

4. Zech, A. et al. Balance training for neuromuscular control and performance enhancement: A systematic review. J. Athl. Train. 45, 392–403 (2010).

5. Billot, M., Simoneau, E. M., Hoecke, J. Van & Martin, A. Age-related relative increases in electromyography activity and torque according to the maximal capacity during upright standing. Eur. J. Appl. Physiol. 109, 669–680 (2010).

6. Riemann, B. L. et al. Comparison of the ankle, knee, hip, and trunk corrective action shown during single-leg stance on firm, foam, and multiaxial surfaces. Arch. Phys. Med. Rehabil. 84, 90–95 (2003).

7. Amiridis, I. G., Hatzitaki, V. & Arabatzi, F. Age-induced modifications of static postural control in humans. Neurosci. Lett. 350, 137–140 (2003).

8. Edwards, A. E., Guven, O., Furman, M. D., Arshad, Q. & Bronstein, A. M. Electroencephalographic Correlates of Continuous Postural Tasks of Increasing Difficulty. Neuroscience 395, 35–48 (2018).

9. Baker, S. N., Olivier, E. & Lemon, R. N. Coherent oscillations in monkey motor cortex and hand muscle EMG show task-dependent modulation. J. Physiol. 501, 225–241 (1997).

10. Conway, B. A. et al. Synchronization between motor cortex and spinal motoneuronal pool during the performance of a maintained motor task in man. J. Physiol. 489, 917–924 (1995).

11. Ushiyama, J. et al. Between-subject variance in the magnitude of corticomuscular coherence during tonic isometric contraction of the tibialis anterior muscle in healthy young adults. J. Neurophysiol. 106, 1379–1388 (2011).

12. Ushiyama, J., Yamada, J., Liu, M. & Ushiba, J. Individual difference in β-band corticomuscular coherence and its relation to force steadiness during isometric voluntary ankle dorsiflexion in healthy humans. Clin. Neurophysiol. 128, 303–311 (2017).

13. Masakado, Y. et al. EEG-EMG coherence changes in postural tasks. Electromyogr. Clin. Neurophysiol. 48, 27–33 (2008).

14. Jacobs, J. V., Wu, G. & Kelly, K. M. M. Evidence for beta corticomuscular coherence during human standing balance: Effects of stance width, vision, and support surface. Neuroscience 298, 1–11 (2015).

15. van Melick, N., Meddeler, B. M., Hoogeboom, T. J., Nijhuis-van der Sanden, M. W. G. & van Cingel, R. E. H. How to determine leg dominance: The agreement between self-reported and observed performance in healthy adults. PLoS One 12, 1–9 (2017).

16. Clark, R. A. et al. Validity and reliability of the Nintendo Wii Balance Board for assessment of standing balance. Gait Posture 31, 307–310 (2010).

17. Clark, R. A., Hunt, M., Bryant, A. L. & Pua, Y. H. Author response to the letter: On ‘ Validity and reliability of the Nintendo Wii Balance Board for assessment of standing balance’[: are the conclusions stated by the authors justified? Gait Posture 39, 1151–1154 (2014).

18. Holmes, J. D., Jenkins, M. E., Johnson, A. M., Hunt, M. A. & Clark, R. A. Validity of the Nintendo Wii® balance board for the assessment of standing balance in Parkinson’s disease. Clin. Rehabil. 27, 361–366 (2013).

19. Imaizumi, S., Asai, T., Hiromitsu, K. & Imamizu, H. Voluntarily controlled but not merely observed visual feedback affects postural sway. PeerJ 6, e4643 (2018).

20. Hjorth, B. An on-line transformation of EEG scalp potentials into orthogonal source derivations. Electroencephalogr. Clin. Neurophysiol. 39, 526–530 (1975).

21. Myers, L. J. et al. Rectification and non-linear pre-processing of EMG signals for cortico-muscular analysis. J. Neurosci. Methods 124, 157–165 (2003).

22. Ward, N. J., Farmer, S. F., Berthouze, L. & Halliday, D. M. Rectification of EMG in low force contractions improves detection of motor unit coherence in the beta-frequency band. J. Neurophysiol. 110, 1744–1750 (2013).

23. Dakin, C. J., Dalton, B. H., Luu, B. L. & Blouin, J. S. Rectification is required to extract oscillatory envelope modulation from surface electromyographic signals. J. Neurophysiol. 112, 1685–1691 (2014).

24. Farmer, S. F., Bremner, F. D., Halliday, D. M., Rosenberg, J. R. & Stephens, J. a. The frequency content of common synaptic inputs to motoneurones studied during voluntary isometric contraction in man. J. Physiol. 470, 127–155 (1993).

25. Gross, J. et al. Cortico[muscular synchronization during isometric muscle contraction in humans as revealed by magnetoencephalography. J. Physiol. 527, 623–631 (2000).

26. Halliday, D. M. et al. A framework for the analysis of mixed time series/point process data—Theory and application to the study of physiological tremor, single motor unit discharges and electromyograms. Prog. Biophys. Mol. Biol. 64, 237–278 (1995).

27. Ushiyama, J., Takahashi, Y. & Ushiba, J. Muscle dependency of corticomuscular coherence in upper and lower limb muscles and training-related alterations in ballet dancers and weightlifters. J. Appl. Physiol. 109, 1086–1095 (2010).

28. Prieto, T. E., Myklebust, J. B., Hoffmann, R. G., Lovett, E. G. & Myklebust, B. M. Measures of postural steadiness: Differences between healthy young and elderly adults. IEEE Trans. Biomed. Eng. 43, 956–966 (2002).

29. Ushiyama, J. & Masani, K. Relation between postural stability and plantar flexors muscle volume in young males. Med. Sci. Sports Exerc. 43, 2089–2094 (2011).

30. Thach, W. T. & Bastian, A. J. Role of the cerebellum in the control and adaptation of gait in health and disease. Prog. Brain Res. 143, 353–366 (2004).

31. Takakusaki, K., Oohinata-sugimoto, J., Saitoh, K. & Habaguchi, T. Role of basal ganglia-brainstem systems in the control of postural muscle tone and locomotion. Prog. Brain Res. 143, 231–237 (2004).

32. Winter, D. Human balance and posture control during standing and walking. Gait Posture 3, 193–214 (1995).

33. Gatev, P., Thomas, S., Kepple, T. & Hallett, M. Feedforward ankle strategy of balance during quiet stance in adults. J. Physiol. 514, 915–928 (1999).

34. Loram, I. D., Maganaris, C. N. & Lakie, M. Human postural sway results from frequent, ballistic bias impulses by soleus and gastrocnemius. J. Physiol. 564, 295–311 (2005).

35. Vieira, T. M. M., Loram, I. D., Muceli, S., Merletti, R. & Farina, D. Recruitment of motor units in the medial gastrocnemius muscle during human quiet standing: is recruitment intermittent? What triggers recruitment? J. Neurophysiol. 107, 666–676 (2012).

36. Sasagawa, S., Ushiyama, J., Masani, K., Kouzaki, M. & Kanehisa, H. Balance control under different passive contributions of the ankle extensors: Quiet standing on inclined surfaces. Exp. Brain Res. 196, 537–544 (2009).

37. Mochizuki, G., Semmler, J. G., Ivanova, T. D. & Garland, S. J. Low-frequency common modulation of soleus motor unit discharge is enhanced during postural control in humans. Exp. Brain Res. 175, 584–595 (2006).

38. Fitzpatrick, B. Y. R. C., Taylor, J. L. & Mccloskey, D. I. Ankle stiffness of standing humans in response to iperceptible perturbation: reflex and task-dependent components. J. Physiol. 533–547 (1992).

39. Watanabe, T., Saito, K., Ishida, K., Tanabe, S. & Nojima, I. Coordination of plantar flexor muscles during bipedal and unipedal stances in young and elderly adults. Exp. Brain Res. 236, 1229–1239 (2018).

40. Héroux, M. E., Dakin, C. J., Luu, B. L., Inglis, J. T. & Blouin, J.-S. S. Absence of lateral gastrocnemius activity and differential motor unit behavior in soleus and medial gastrocnemius during standing balance. J. Appl. Physiol. 116, 140–148 (2014).

41. Lemos, T., Imbiriba, L. A., Vargas, C. D. & Vieira, T. M. Modulation of tibialis anterior muscle activity changes with upright stance width. J. Electromyogr. Kinesiol. 25, 168–174 (2015).

42. Sozzi, S., Honeine, J. L., Do, M. C. & Schieppati, M. Leg muscle activity during tandem stance and the control of body balance in the frontal plane. Clin. Neurophysiol. 124, 1175–1186 (2013).

43. Hwang, S. et al. The balance recovery mechanisms against unexpected forward perturbation. Ann. Biomed. Eng. 37, 1629–1637 (2009).

44. Gwin, J. T. & Ferris, D. P. Beta- and gamma-range human lower limb corticomuscular coherence. Front. Hum. Neurosci. 6, 1–6 (2012).

45. Omlor, W., Patino, L., Hepp-Reymond, M. C. & Kristeva, R. Gamma-range corticomuscular coherence during dynamic force output. Neuroimage 34, 1191–1198 (2007).

46. Brown, P. et al. Cortical correlate of the piper rhythm in humans. J. Neurophysiol. 80, 2911–2917 (1998).

47. Ushiyama, J. et al. Contraction level-related modulation of corticomuscular coherence differs between the tibialis anterior and soleus muscles in humans. J. Appl. Physiol. 112, 1258–1267 (2012).

48. Mima, T., Simpkins, N., Oluwatimilehin, T. & Hallett, M. Force level modulates human cortical oscillatory activities. Neurosci. Lett. 275, 77–80 (1999).

49. Nunez, P. L. Generation of human EEG by a combination of long and short range neocortical interactions. Brain Topogr. 1, 199–215 (1989).

50. Yan, Y., Nunez, P. L. & Hart, R. T. Finite-element model of the human head: scalp potentials due to dipole sources. Med. Biol. Eng. Comput. 29, 475–481 (1991).

51. Malmivuo, J., Suihko, V. & Eskola, H. Sensitivity distributions of EEG and MEG measurements. IEEE Trans. Biomed. Eng. 44, 196–208 (1997).

52. Farina, D., Cescon, C. & Merletti, R. Influence of anatomical, physical, and detection-system parameters on surface EMG. Biol. Cybern. 86, 445–456 (2002).

53. Mesin, L., Merletti, R. & Rainoldi, A. Surface EMG: The issue of electrode location. J. Electromyogr. Kinesiol. 19, 719–726 (2009).

